# Highly prevalent co-infections and genetic diversity of *Anaplasma* species in asymptomatic small ruminants in a Mediterranean context revealed by a *gltA-*based nested multiplex PCR assay

**DOI:** 10.64898/2025.12.17.695055

**Authors:** Maggy Jouglin, Léana Rodrigues, Claire Bonsergent, Suzanne Bastian, Laurence Malandrin

## Abstract

Anaplasmosis is a widespread tick-borne disease affecting various host species. Small ruminants (sheep and goats) can be infected by multiple *Anaplasma* species:. To evaluate the circulation and co-infection rates of five *Anaplasma* species (*A. ovis*, *A. capra*, *A. bovis*, *A. marginale* and *A. phagocytophilum*) in asymptomatic small ruminants in Corsica—a Mediterranean island of France—we designed a singleplex, duplex, and multiplex nested PCR (nPCR) scheme based on the *gltA* gene. Anaplasmosis was found to be highly endemic on the island, with 85.2% of the tested small ruminants infected by one or more *Anaplasma* species (82.8% of sheep and 89.5% of goats). We confirmed the presence of *A. ovis* and *A. capra*, and, for the first time in Corsica, detected *A. bovis* and *A. phagocytophilum* in both sheep (respective infection rates: 64.3%, 34.3%, 4.3%, and 7.1%) and goats (respective rates: 84.2%, 13.2%, 18.4%, and 5.3%). *A. marginale* was not detected. Approximately one-third of the infected animals were co-infected (35.1% of infected sheep and 32.4% of infected goats). *A. ovis* was always involved in co-infections, most often associated with *A. capra* (54.8% of co-infections). Importantly, *A. bovis* was never detected as a mono-infection in asymptomatic animals, but was always found associated with *A. ovis* (32.3% of co-infections). Sequence analysis of the *gltA* and 16S rRNA genes confirmed the species identification of all detected *Anaplasma* species. A probably new *Anaplasma* species was detected for the first time in Corsica using the 16S rRNA nPCR. Furthermore, we found several circulating *A. phagocytophilum* genetic variants. Sequence analysis of *A. ovis* 16S rRNA revealed a single SNP that delineates two sequence groups (Ao1goat and Ao2sheep), which were significantly linked to the host of origin. Additionally, two groups of *A. bovis gltA* sequences were identified (Ab1 and Ab2, with 87.4% sequence homology), forming two novel clades. While the Ab1 clade is closely related to the six previously described clades, the Ab2 clade formed a separate phylogenetic group that also contained isolates from Corsican symptomatic goats. In conclusion, *Anaplasma* species are highly prevalent in the small ruminant population of Corsica, confronted with a great diversity of co-infecting species and genetic variants.

## 1. Introduction

Rickettsial alphaproteobacteria of the genus *Anaplasma* are obligate intracellular bacteria that can infect a broad range of vertebrate species and are transmitted by ticks. At present, 8 species and several unclassified genovariants have been described, however, some species delineations are still under debate (Rar et al., 2021). Different species of *Anaplasma* have the ability to infect different hosts and different kinds of blood cells. They may be detrimental to the hosts through fever, anemia or interference with the immune system (Atif et al., 2016).

Most *Anaplasma* species have a broad host range, a characteristic particularly evident for *A. phagocytophilum*, *A. bovis*, and more recently described for *A. capra* (Atif, 2016; Altay et al., 2024). Given that these bacteria can establish persistent infections, the occurrence of co-infections with multiple *Anaplasma* species may not be uncommon and may significantly impact animal health. Small ruminants, including both sheep and goats, are frequently susceptible to infection by at least four *Anaplasma* species: *A. ovis*, *A. capra*, *A. bovis* and *A. phagocytophilum* (Atif et al., 2016).

For instance, on the Mediterranean island of Corsica, France, the co-circulation of *A. capra* and *A. ovis* in goat has been independently reported in recent publications (Cabezas-Cruz et al., 2019; Jouglin et al., 2022). However, these studies did not specifically investigate the presence of co-infections. In contrast, a recent study in Türkiye highlighted the exclusive occurrence of *A. ovis*, with no detection of *A. bovis*, *A. capra*, *A. platys*-like, or *A. phagocytophilum* (Karatepe et al., 2025). Recognizing the confirmed presence of at least two *Anaplasma* species in small ruminants in Corsica, we aimed to evaluate the possibility of co-infections with multiple *Anaplasma* species using existing sheep and goat samples that had previously only been analyzed for *A. capra*.

To achieve this goal, we developed a novel PCR assay. Various gene targets are commonly employed for detecting and analyzing the genetic diversity of *Anaplasma* species (Rar et al., 2021). Among these, some genetic markers, such as *ankA* for *A. phagocytophilum* (Sharf et al., 2011), are highly species-specific. Others, like the *msp* genes, are excessively variable for a multi-species PCR approach, while highly conserved genes, such as the 16S rRNA gene, offer insufficient resolution for our specific purpose.

The two constitutive genes, *groEL* and *gltA*, are also frequently utilized to assess the genetic diversity within several *Anaplasmataceae* species. Studies using these markers often reveal the existence of distinct clusters and sub-population structures within certain species (Bauer et al., 2021; Jouglin et al., 2022). The widespread use of these common molecular markers has led to a rich repository of comparable sequences in databases, derived from diverse bacterial species and genotyped from a broad range of hosts and geographical locations. Between the two, *gltA* has been reported to be more variable than to *groEL* when both sequences from the same isolates were obtained (Aung et al., 2024). Following a preliminary sequence comparison of both gene targets, *gltA* was selected for the current study. These characteristics not only facilitate the development of a multiplex assay but also enable a comprehensive study of the genetic diversity within the different *Anaplasma* species.

To investigate the extent and diversity of co-infections in small ruminants, we developed a multiplex PCR protocol. This assay, based on the genetic diversity of the *gltA* gene, allows for the comprehensive evaluation of co-infections by *A. ovis, A. capra, A. bovis, A. phagocytophilum*, and *A. marginale*.

## 2. Materials and methods

### 2.1. Sample populations

Blood and/or DNA samples collected during a previous study in Corsica, France (Jouglin et al., 2022), were re-utilized for this investigation. These samples were originally collected in 2020 to screen for *Anaplasma capra*. For the current study, a multiplex nested PCR was developed to assess co-infections with various *Anaplasma* species in sheep and goats from Corsica.

Briefly, samples were obtained from eleven farms, comprising 70 asymptomatic sheep and 38 asymptomatic goats (Figure 1). Additionally, four blood samples from symptomatic goats of unknown localization in Corsica and sent for *Anaplasma* spp. diagnosis were included. Specific details regarding the symptoms are unavailable as these samples were submitted for diagnostic purposes without further clinical information. All samples and DNA were prepared as previously described (Jouglin et al., 2017).

**Fig. 1.**
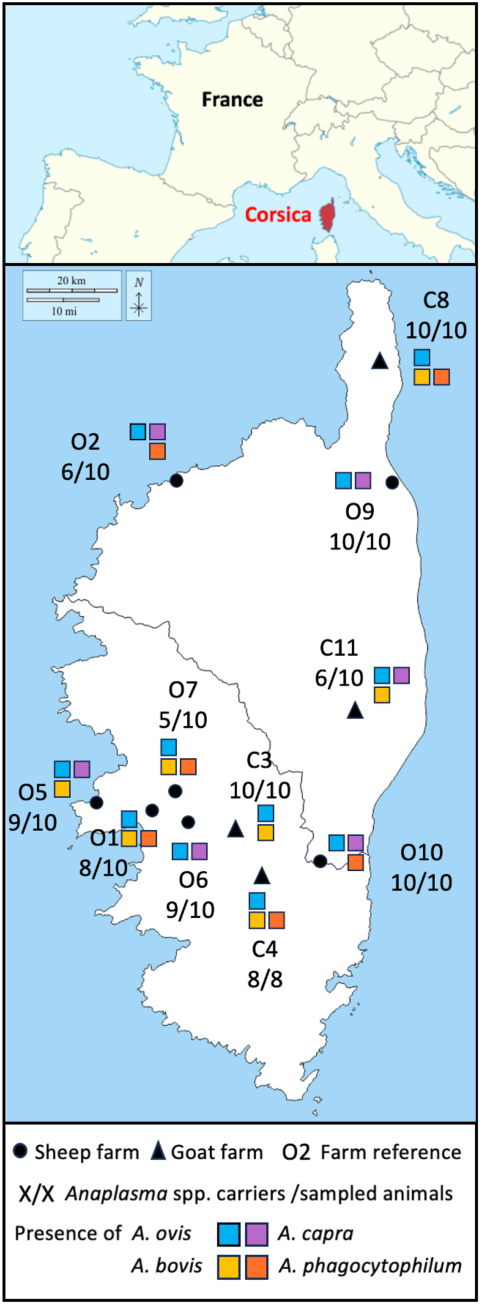
Localization of Corsica in Europe, and of the ovine and caprine farms in Corsica. The number of infected animals as well as the identities of the detected *Anaplasma* spp. are indicated for each farm.

### 2.2. Multiplex PCR Assay Design

To develop a species-specific multiplex PCR, the *gltA* gene was selected due to its constitutive nature, genetic diversity, and the availability of diverse isolate sequences in public databases at the time of study design (2020). Representative *gltA* sequences were chosen for their broad host range and geographical origins. A comprehensive list of these sequences for each target species (*A. capra, A. ovis, A. bovis, A. phagocytophilum*, and *A. marginale*) is provided in Supplementary Table 1.

These sequences were aligned using Clustal Omega Multiple Sequence Alignment (MSA) (Sievers et al., 2011) to generate a consensus *gltA* sequence for each species. Then regions conserved within each species but distinct between species were searched for. The goal was to design primers that would amplify targets at similar melting temperatures while producing amplicons of varying sizes, allowing for their differentiation on agarose gels. Given that *Anaplasma* bacteremia can be low in asymptomatic animals, a nested multiplex PCR approach was implemented to enhance sensitivity.

Based on these consensus alignments, we concluded that a single-step multiplex nested PCR would not be feasible. Therefore, we designed two distinct first-round PCRs: one for *A. bovis* and *A. phagocytophilum*, and another for *A. capra, A. ovis*, and *A. marginale*. These separate first-round amplification products were then mixed in equal volumes and diluted (1:50) for a single second-round PCR using species-specific primers (Fig. 2).

**Fig. 2.**
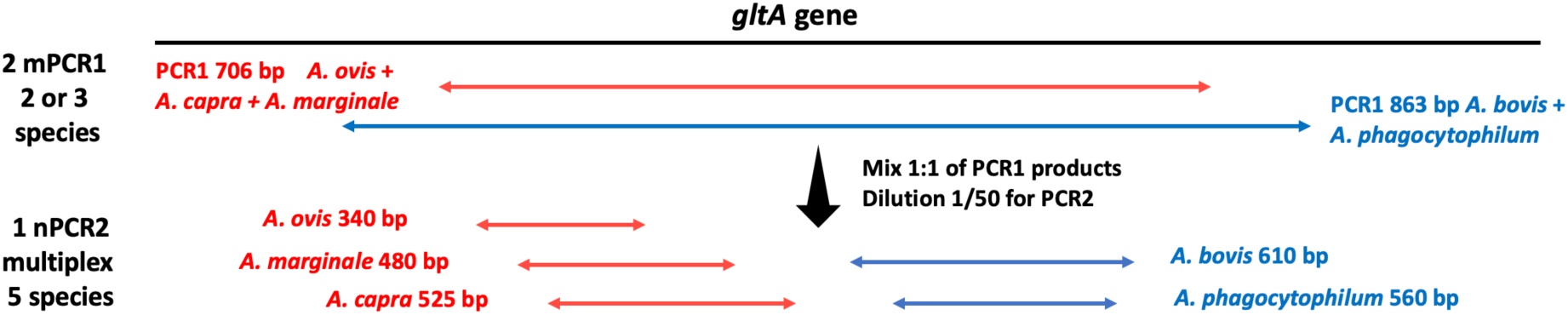
Schematic representation of the nested mPCR amplification protocols used in this study.

The specificity of each primer set was evaluated in silico using the Basic Local Alignment Search Tool (BLAST: https://blast.ncbi.nlm.nih.gov/Blast.cgi) before their evaluation on samples in singleplex and multiplex nPCR. The complete multiplex nested PCR design and the specific primers selected are detailed in Figure 2 and Table 1, respectively.

**Table 1.**
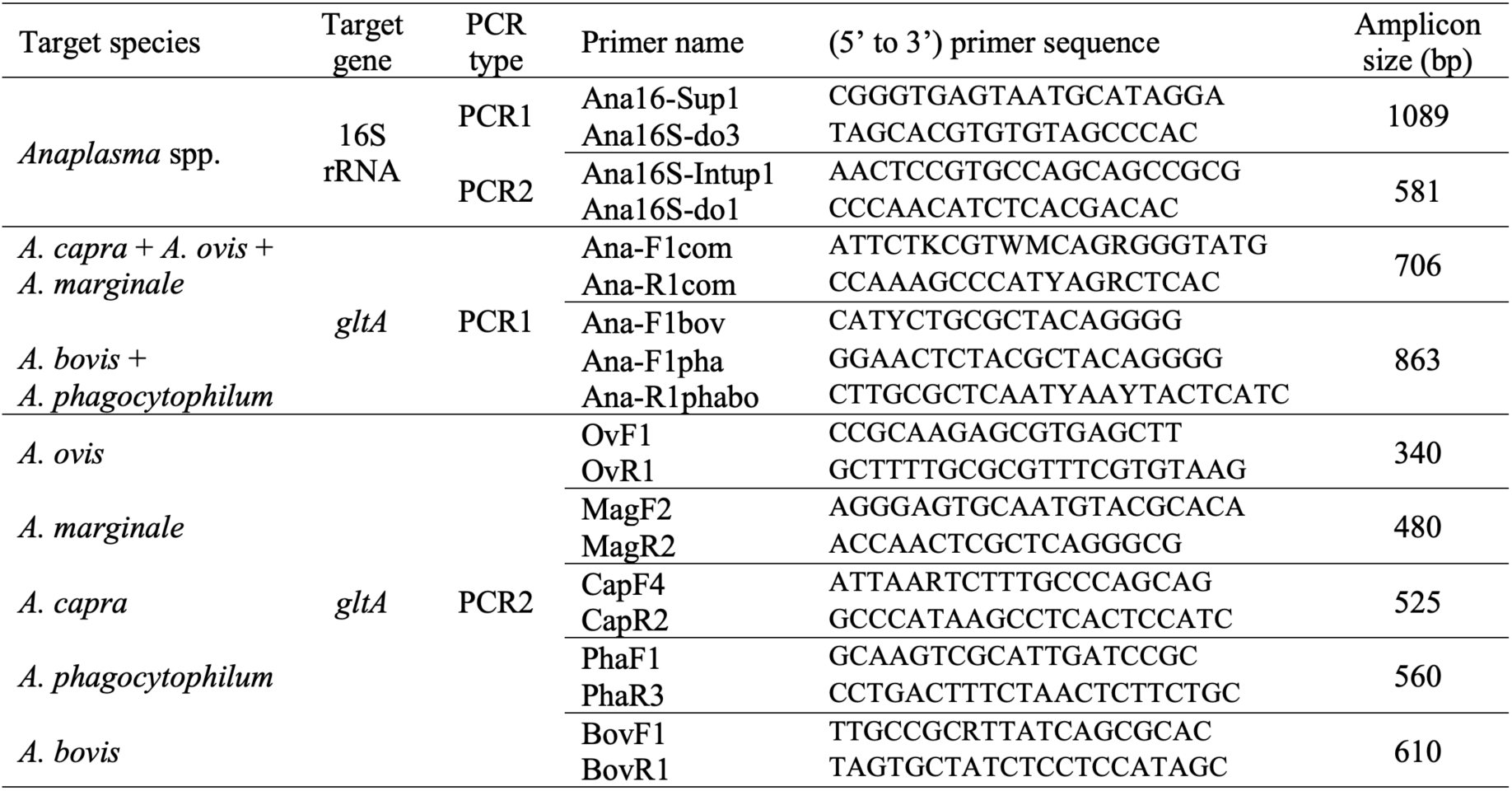
List of primers used for the detection and characterization of *Anaplasma* species.

### 2.3. Origin of *Anaplasma* spp. positive samples and DNA

All 112 sheep and goat samples from a previous study (Jouglin et al., 2022) were initially screened for *Anaplasma* spp. using a previously described nested PCR targeting the 16S rRNA ribosomal gene (Jouglin et al., 2019). All positive amplicons (approximately 500 bp) were sequenced and submitted to a BLASTn search to identify the *Anaplasma* species detected.

*Anaplasma capra, A. ovis*, and *A. bovis*-infected samples were selected from the goat and sheep blood samples tested as described above. Cervid blood found to be infected with *A. phagocytophilum* was sourced from a previous study conducted in a wild fauna reserve in France (Jouglin et al., 2019). Finally, *Anaplasma marginale* DNA from infected sheep was a kind gift from Professor Furhan Iqbal (Bahauddin Zakariya University, Pakistan), originating from a previous study (Abid et al., 2021).

DNA samples confirmed positive for *Anaplasma spp.* were then used to amplify *gltA* gene fragments. These fragments were subsequently cloned and utilized both to evaluate the detection limit of our assays and as positive controls in single and multiplex PCR reactions.

### 2.4. Amplification and cloning of partial *gltA* genes

Partial *gltA* genes from each target *Anaplasma* species were amplified and subsequently cloned to generate single-species DNA controls. These controls were used to assess the detection threshold for each *Anaplasma* species. Primers selected for these amplifications were listed in table 1.

Amplification was performed using Pfu DNA polymerase (Promega, USA), which possesses proofreading activity to minimize amplification errors. Amplicons were purified using a PCR clean-up kit (Macherey Nagel, Germany), then ligated into the pGEM®-T Easy plasmid vector (Promega, USA). The ligated plasmids were subsequently cloned by transformation into JM 109 *Escherichia coli* cells, following the manufacturer’s instructions. To confirm the successful insertion of the expected gene fragments, plasmid DNA was extracted using NucleoSpin Plasmid (Macherey Nagel, Germany), quantified with the Qubit™ dsDNA HS Assay kit (Invitrogen), and sequenced bi-directionally (GATC, Germany) using the plasmid primers T7 and SP6.

### 2.5. Optimization of Multiplex PCR

We first determined the optimal PCR amplification conditions, including DNA concentration and annealing temperature, for each primer set, as well as sensitivity. This was done using calibrated cloned *gltA* fragments, prepared as tenfold serial dilutions ranging from 10^6^ to 10^0^ copies of plasmid per microliter. For the five primer sets used in the multiplex second-round PCR, the optimum and common annealing temperature was selected using a temperature gradient from 50°C to 60°C. For overall optimization, we tested various concentrations of dNTPs (0.2 mM to 0.45 mM), MgCl_2_ (2 mM to 3 mM), and primers (0.4 μM to 1 μM).

Once the PCR conditions were adjusted to allow efficient amplification of each species’ *gltA* fragment, we further tested specificity by progressively running multiplex PCRs. This began with duplex PCRs, leading up to multiplex PCRs incorporating all five primer sets.

The optimized PCR conditions were as follows. For the two PCR1, 5 μl of genomic DNA served as the template in a final volume of 30 μl. The reaction mixture contained 0.45 mM dNTPs (Eurobio, France), 1X PCR buffer, 3 mM MgCl2, 1 μM of each primer (*A. capra*, *A. marginale, A. ovis* in one reaction, *A. phagocytophilum* and *A. bovis* in the other one), 0.15 μl (5 U/μl) of GoTaq Flexi (Promega, USA), and deionized water. The PCR protocol included an initial denaturation step at 95°C for 3 min, followed by 40 cycles of 30 s at 95°C, 30 s at 56°C, and 30 s at 72°C. A final elongation step of 5 min at 72°C concluded the reaction.

The nested PCRs were then performed using 10 μl of the first-round amplicons (diluted 1:50 of an 1/1 ratio of each first PCR amplicon) as a template (Fig. 2). The PCR2 was performed in the same conditions as PCR1 except the primers concentrations (0.84 μM of each primer for *A. ovis* and 1.05 μM of each primer for *A capra*, *A. marginale, A. bovis* and *A. phagocytophilum)*.

Two negative controls were included in each experiment: an extraction control and a PCR contamination control (blank). Amplicons were visualized and their sizes verified under UV light on a 1.5% agarose gel stained with ethidium bromide.

### 2.6. Sequencing and phylogenetic analysis

To confirm the specific amplification of each target species, single amplicons were purified using ExoSAP-IT (Ozyme, France) and sent for conventional Sanger sequencing in both directions (GATC, Germany).

For the phylogentic analysis, the longest and representative sequences were selected and were trimmed to generate *gltA* partial sequences ranging from 415 to 420 bp for *A. bovis* and of 500 bp for *Anaplasma* spp. 16S rRNA sequences. The two trees were built with respectively, published sequences that represent different *A. bovis* variants, and various 16S rRNA sequences of *Anaplasma* species relevant to our study. *Ehrlichia canis* sequences of the *gltA* gene and 16S rRNA gene were used as an outgroup.

For both the *A. bovis* partial *gltA* sequences and the *Anaplasma* spp. 16S rRNA phylogenetic analyses, Maximum Likelihood phylogenetic trees were constructed using the FastTree workflow provided by the NGPhylogeny platform (https://ngphylogeny.fr)(Lemoine et al., 2019). The alignments were performed using MAFFT software (Katoh and Standley, 2013), cleaned with BMGE software (Criscuolo and Gribaldo, 2010) and phylogenetic trees were built using FastTree software (Price et al., 2009; Price et al., 2010). The final trees were generated in Newick format (Junier and Zdobnov, 2010). The GTR evolutionary model was used. For both trees, bootstrapping support values for the topology have been calculated with 1000 replicates (Lemoine et al., 2018).

## 3. Results

### 3.1. Multiplex PCR design, specificity and species detection lower limit of detection

For each species, the primers selected in silico to specifically amplify the 5 *Anaplasma* species were tested first on the plasmid extracted DNA containing the corresponding *gltA* inserts, and amplicons of the expected sizes were obtained: *A. ovis* 340 bp, *A. marginale* 480 bp, *A. capra* 525 bp, *A. phagocytophilum* 560 bp and *A. bovis* 610 bp. The primers were then tested with plasmid extracted DNA of each of the other species either alone or as a mixture, and amplifications were observed only with the species corresponding DNA (data not shown), confirming primer specific amplifications. The detection threshold of each PCR was determined using plasmids containing partial *gltA* genes of each tested species. The detection limits for *A. bovis* and *A. capra* were between 1 and 10 copies per reaction and *A. phagocytophilum*, *A. marginale* and *A. ovis* between 10 and 100 copies per reaction.

Then each DNA extracted from blood samples was analyzed using 16S rRNA nPCR, singleplex, duplex (*A. ovis* + *A. capra* and *A. bovis* + *A. phagocytophilum*) and multiplex *gltA* nPCR (*A. ovis*, *A. capra*, *A. bovis* and *A. phagocytophilum*)(Fig. 3). The 16S rRNA amplicons were sent for sequencing as species identification controls (sequence analysis is detailed in section 3.6). Similarly, some single *gltA* amplicons were sent for sequencing.

**Fig. 3.**
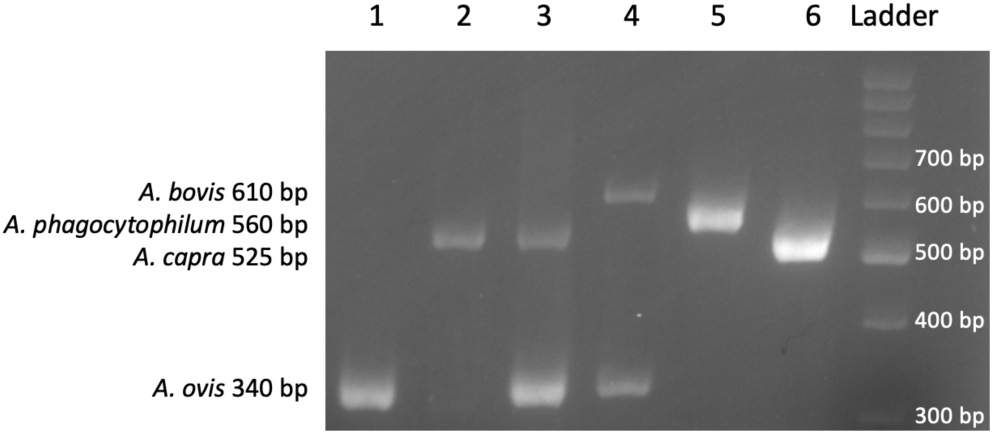
Evidence of *Anaplasma* spp. co-infections in small ruminants’ blood samples. 1 = *A. ovis* in sheep, 2 = *A. capra* in sheep, 3 = coinfection of *A. ovis*/*A. capra* in sheep, 4 = coinfection of *A. ovis*/*A. bovis* in goat, 5 = *A. phagocytophilum* in sheep, 6 = *A. capra* in goat.

Five *Anaplasma* species were detected in the 112 samples: *A. ovis*, *A. capra*, *A. bovis, A. phagocytophilum* and *Anaplasma* sp., the latter not being included in our multiplex detection scheme. As *A. marginale* was not detected using *gltA*-based singleplex PCR, its search in duplex and multiplex PCR was not performed.

### 3.2. Comparison of detection with nested 16S rRNA nPCR and *gltA*-based singleplex, duplex PCR or multiplex nPCR on asymptomatic small ruminants

The detection efficiency and reliability of each PCR method were then compared on the 108 blood samples of asymptomatic small ruminants (table 2, figure 4 and supplementary table 2).

**Fig. 4.**
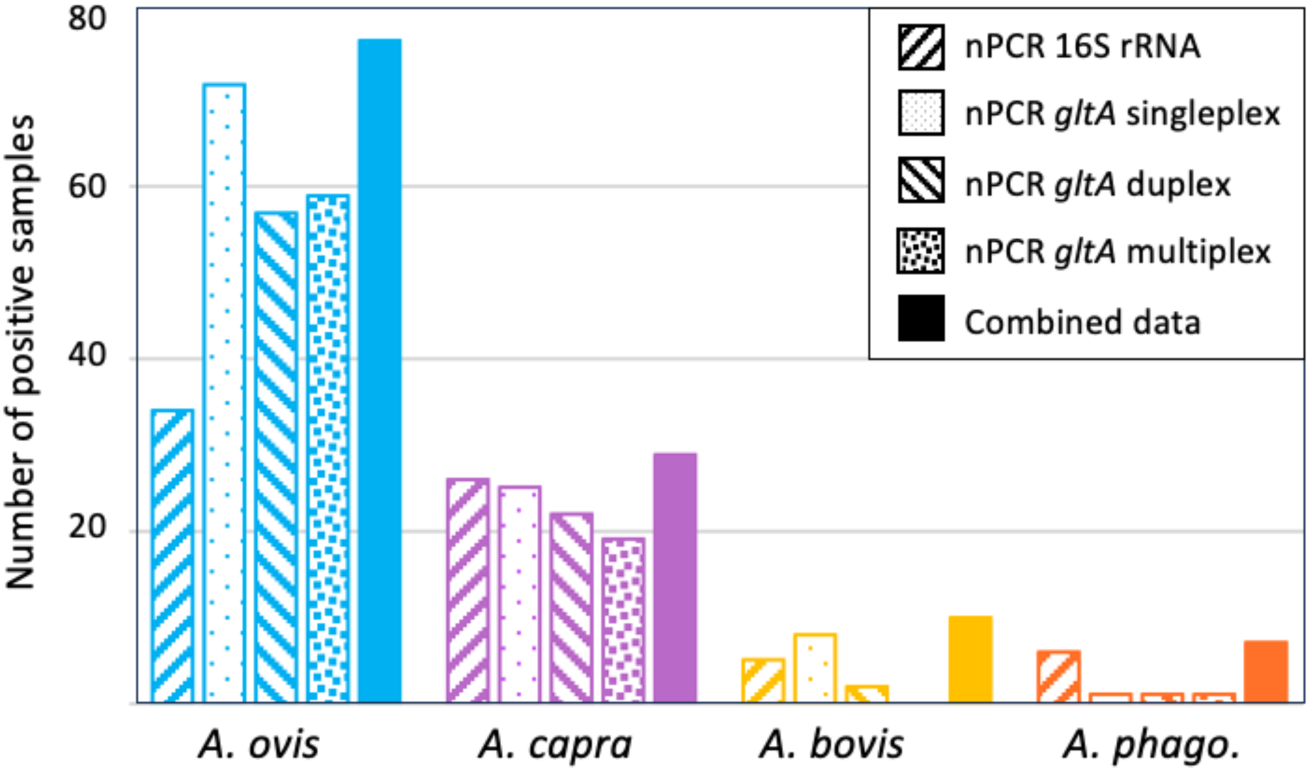
Detection of *Anaplasma* spp. in asymptomatic small ruminants in Corsica according to the method used (16S rRNA nPCR, singleplex, duplex or multiplex *gltA* nPCR).

**Table 2.**
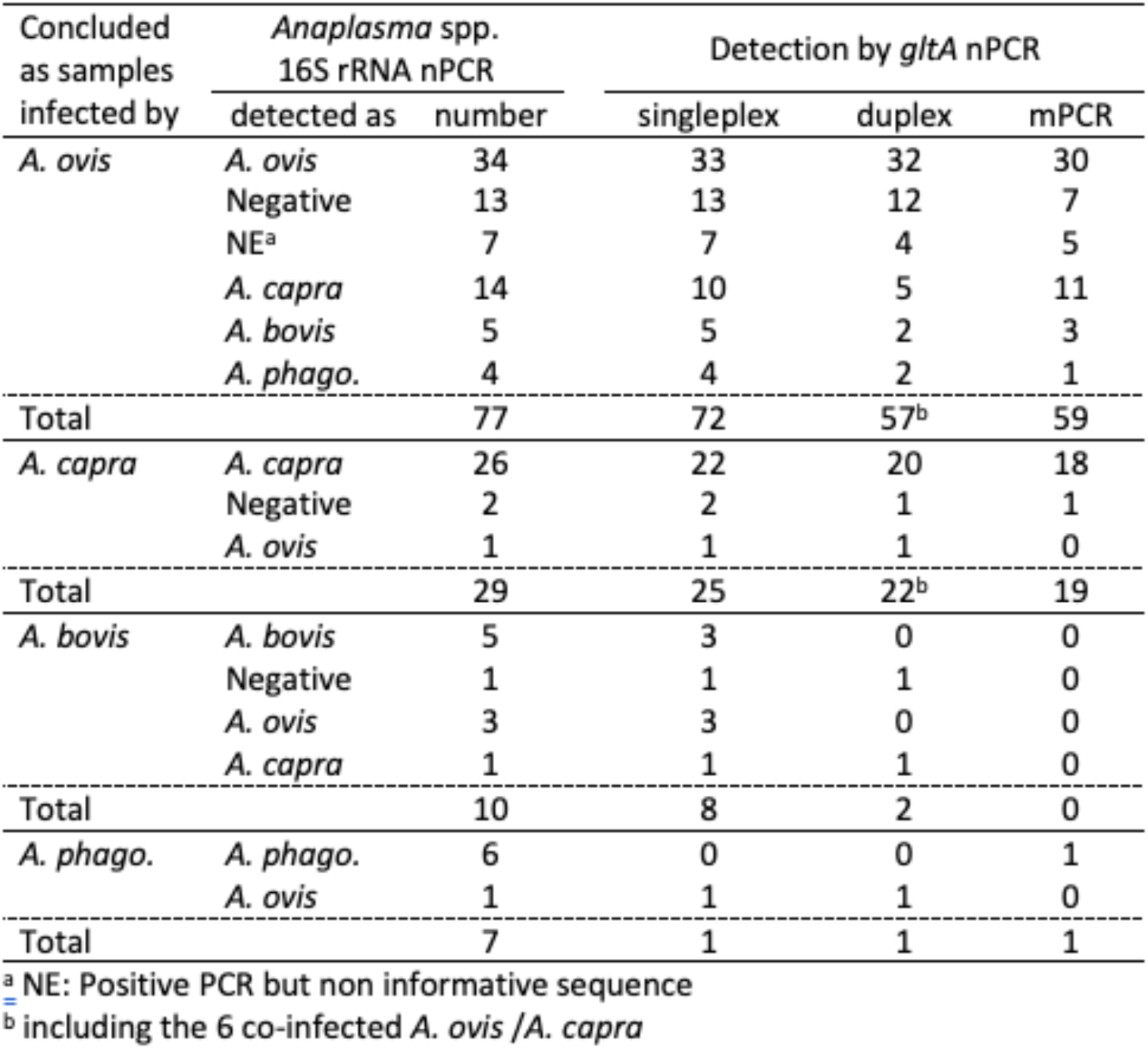
Comparison of the detection of *Anaplasma ovis*, *A. capra*, *A. bovis* and *A. phagocytophilum* (*A. phago*.) in asymptomatic small ruminants’ blood samples form Corsica (France) according to the detection methods used. For example, among the 13 samples that were not detected using the 16S rRNA nPCR (Negative), *A. ovis* was detected in all of them using the singleplex *gltA*-based nPCR, in 12/13 using the duplex nPCR *A. ovis*/*A. capra* and in 7/13 using the multiplex nPCR. Another example: among 14 samples identified by PCR and sequencing as mono-infected with *A. capra* with the 16S rRNA nPCR, co-infection with *A. ovis* was found for ten using the *A. ovis* singleplex nPCR, five using the duplex nPCR *A. ovis*/*A. capra* and eleven using the multiplex nPCR.

With the 16S rRNA nPCR, one *Anaplasma* species was detected in 82/108 samples (75.9%). Sequencing was then necessary and species identification was achieved by a BLASTn search in 72 cases: *A. ovis* (34), *A. capra* (26), *A. phagocytophilum* (6) and *A. bovis* (5). One sheep sample was identified as being infected by an *Anaplasma* closely related to *A. platys* and is refered to as *Anaplasma* sp. An *Anaplasma* species could not be attributed to 10 positive samples due to uninterpretable sequences (noted NE in the supplementary table 2).

The specific singleplex sets of *gltA*-based nPCRs managed to detect and identify *Anaplasma* species in 58/72 of the 16S rRNA already identified species (33/34 *A. ovis*, 22/26 *A. capra*, 3/5 *A. bovis* but none of the *A. phagocytophilum*) as well as in 16/26 of the 16S rRNA negative samples (13 *A. ovis*, two *A. capra*, and one *A. bovis*) and in 7/10 infected samples with non-identified *Anaplasma* species (six *A. ovis* and one *A. ovis/A. bovis*). The 5 singleplex sets of nPCRs achieved separatly the detection and direct identification of a total of 81/108 (75%) samples. The *A. bovis* and *A. phagocytophilum* singleplex nPCRs were not very efficient, detecting respectively only 3/6 and 0/6 of the samples tested positive by the 16S rRNA nPCR. They however detected 5 more samples infected with *A. bovis* and one infected by *A. phagocytophilum* (individual results are detailed in Supplementary table 2).

The two separate sets of duplex nPCRs (*A. ovis*/*A. capra* and *A. bovis/A. phagocytophilum*) detected *Anaplasma* spp. in *7*6/108 (70.4%) already identified samples, either as mono-infections (51 *A. ovis*, 16 *A. capra*, two *A. bovis,* one *A. phagocytophilum*) or as co-infections (six *A. capra*/*A. ovis*)(Supplementary table 2). The combined results from the singleplex PCR detected 10 co-infections *A. ovis*/*A. capra*, while only six of them were detected using the duplex PCR. In the four discordant cases, either *A. capra* (2 samples) or *A. ovis* (2 samples) was detected. The second duplex PCR (*A. bovis/A. phagocytophilum/A. marginale)* allowed the detection of 2 samples infected with *A. bovis* and one infected with *A. phagocytophilum*. With the multiplex nPCR, *Anaplasma* spp. was detected in 67/108 samples (62%), either as mono-infections (47 *A. ovis* and eight *A. capra*) or as co-infections (11 *A. capra*/*A. ovis* and one *A. ovis*/*A. phagocytophilum*)(Supplementary table 2). *Anaplasma bovis* was never detected.

### 3.3. Infection rates of *Anaplasma* spp. in asymptomatic sheep and goat in Corsica

We used the combination of the different detection methods (16S rRNA nPCR, *gltA* singleplex, duplex PCRs or the multiplex nPCR) to determine the infection rates by each of the five detected species in the farms, *A. ovis*, *A. capra*, *A. bovis*, *A. phagocytophilum* and *Anaplasma* sp. (Table 3, Figs. 1 and 4).

**Table 3.** Occurrence and infection rates of *Anaplasma* species in asymptomatic small ruminants (sheep and goat) in Corsica as determined by the combined results of the 16S rRNA nPCR, and the *gltA*-based singleplex, duplex and multiplex nPCR.

| Farm reference | Sampled animals | *Anaplasma* spp. | *A. ovis* | *A. capra* | *A. bovis* | *A. phago.* | *Anaplasma* sp. |
| --- | --- | --- | --- | --- | --- | --- | --- |
| O1 | 10 | 8 | 8 | 0 | 1 | 1 | 0 |
| O2 | 10 | 6 | 5 | 1 | 0 | 2 | 0 |
| O5 | 10 | 9 | 9 | 3 | 1 | 0 | 0 |
| O6 | 10 | 9 | 9 | 1 | 0 | 0 | 0 |
| O7 | 10 | 6 | 5 | 0 | 1 | 1 | 1 |
| O9 | 10 | 10 | 6 | 10 | 0 | 0 | 0 |
| O10 | 10 | 10 | 3 | 9 | 0 | 1 | 0 |
| **Total ovine** | **70** | **58** | **45** | **24\*** | **3** | **5** | **1** |
| **% infected** |  | **82.8%** | **64.3%** | **34.3%** | **4.3%** | **7.1%** | **1.4%** |
| C3 | 10 | 10 | 10 | 0 | 3 | 0 | 0 |
| C4 | 8 | 8 | 8 | 0 | 1 | 1 | 0 |
| C8 | 10 | 10 | 10 | 0 | 1 | 1 | 0 |
| C11 | 10 | 6 | 4 | 5 | 2 | 0 | 0 |
| **Total caprine** | **38** | **34** | **32\*** | **5** | **7\*** | **2** | **0** |
| **% infected** |  | **89.5%** | **84.2%** | **13.2%** | **18.4%** | **5.3%** |  |
| **Total small ruminants** | **108** | **92** | **77** | **29** | **10** | **7** | **1** |
| **% infected** |  | **85.2%** | **71.3%** | **26.8%** | **9.3%** | **6.5%** | **0.9%** |

\* significantly different between the sheep and the goat infected populations (Fisher exact test $p < 0.05$ ).

Most of the sampled asymptomatic small ruminants (92/108, 85.2%) were infected by at least one of the *Anaplasma* species we searched for, and no significant difference was found between infection rates of sheep (57/70, 81.4%) or goat (34/38, 89.5%).

*Anaplasma ovis* was the predominant species circulating in small ruminants in Corsica (77/108, 71.3%). It was detected in all sheep or goat farms, with infection rates ranging from 30 to 90% in sheep flocks (mean 64.3%) and from 40 to 100% in goat herds (84.2%), and significantly higher in goat compared to sheep (Fisher’s exact test 0.0439).

*Anaplasma capra* was detected in more sheep farms (5/7) compared to the goat farms (1/4), resulting in an infection rate significantly higher for sheep (34.3%) compared to goat (13.2%)(Fisher’s exact test 0.0227).

*Anaplasma bovis* was detected at least once in the sampled animals of all goat farms resulting in an infection rate of 18.4% significantly higher than the infection rate in sheep (4.3%)(Fisher’s exact test 0.0314), with only one sampled sheep infected in three of the seven farms.

*Anaplasma phagocytophilum* infection rate is rather low in small ruminants (6.5%), with no significant differences between sheep (7.1%) and goat (5.3%). Despite this low infection rate, it was detected in 6/11 farms (4/7 sheep flocks and 2/4 goat herds).

In conclusion, the distribution of the four detected *Anaplasma* species differed between sheep and goat as shown in Fig. 5.

**Fig. 5.**
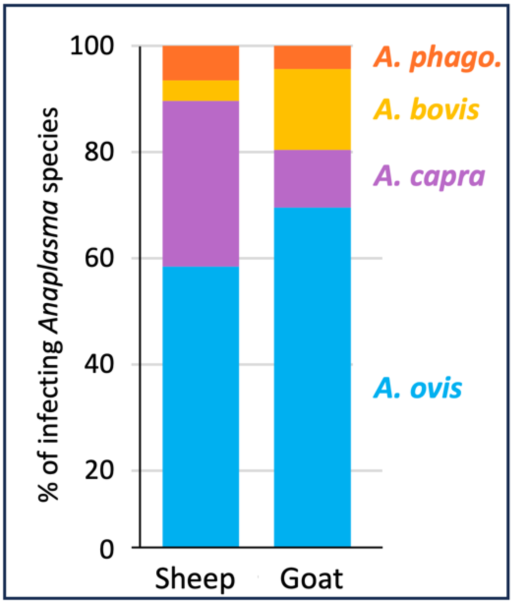
Proportion of the four detected *Anaplasma* species in sheep and goat in Corsica (*A. phago* : *A. phagocytophilum*).

### 3.4. Evaluation of *Anaplasma* spp. co-infections in asymptomatic sheep and goat in Corsica

This study highlighted the high level of co-infections by the four *Anaplasma* species in small ruminants. Among the infected small ruminant population analyzed, 20 sheep (35.1%) and 11 goat (32.4%) were co-infected by at least two different *Anaplasma* species (Fig. 6).

**Fig. 6.**
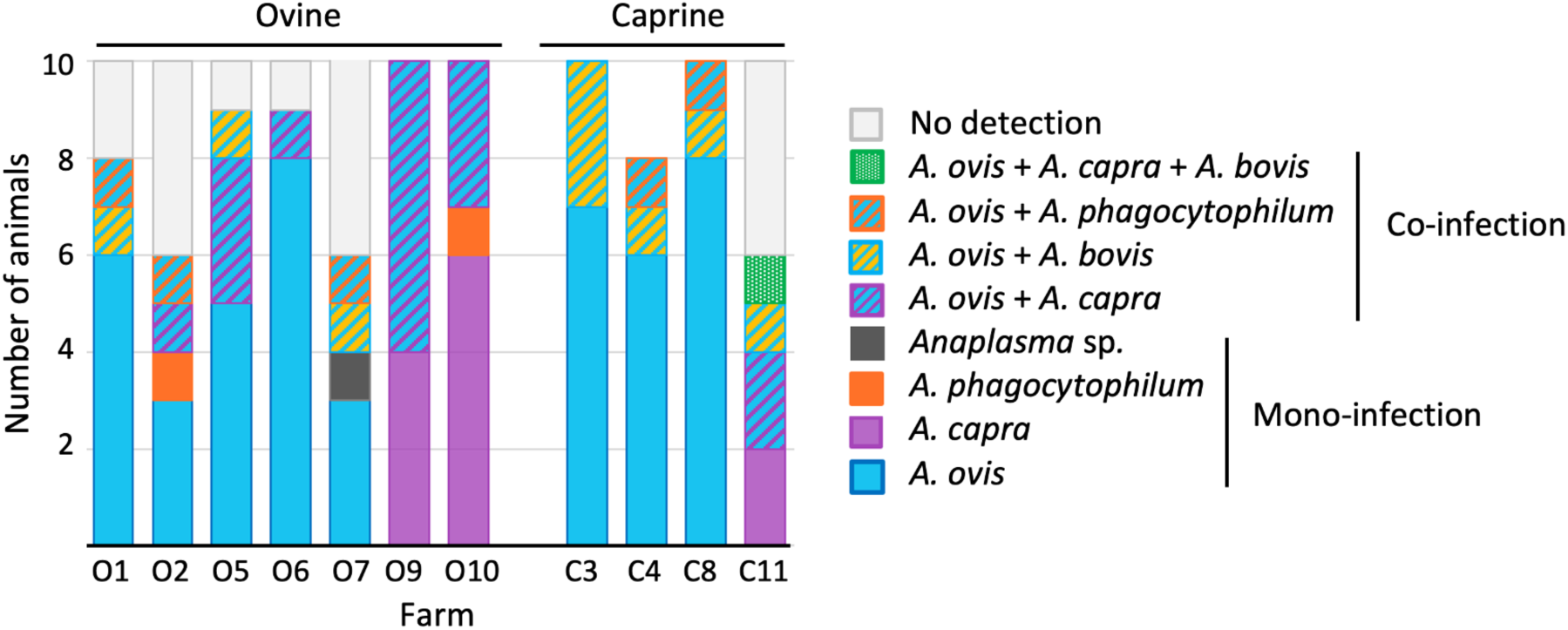
Infection rates, identities and co-infections with five *Anaplasma* species in sheep and goat farms in Corsica. The data corresponds to the combined detection of *Anaplasma* spp. with either 16S rRNA nPCR, or *gltA* singleplex, duplex and multiplex nPCR analyses.

All co-infections involved *A. ovis*, associated with either *A. capra* (17/31, 54.8%), *A. bovis* (10/31, 32.3%) or *A. phagocytophilum* (5/31, 16.1%). A triple co-infection was evidenced only once in a goat involving *A. ovis*, *A. capra* and *A. bovis*.

Co-infections with *A. ovis* and *A. capra* are significantly more frequent in sheep (14/20, 70%) compared to goats (2/11, 18.2%)(Fisher’s exact test 0.0091), while co-infections with *A. ovis* and *A. bovis* are on the contrary significantly more frequent in goats (7/11, 63.4%) compared to sheep (3/20, 15%)(Fisher’s exact test 0.0134). Three *A. ovis* infected sheep (15% of the co-infected animals) and two goats (18.2%) were co-infected with *A. phagocytophilum* (not significantly different).

*Anaplasma bovis* was never detected in a mono-infected small ruminant, but was always co-infecting with *A. ovis* (three sheep and seven goats).

### 3.5. Detection of *Anaplasma* spp. in symptomatic goat

Four blood samples from symptomatic goats were sent by a routine diagnosis laboratory from Corsica, for confirming the presence of *Anaplasma,* without details about symptoms or the precise location on the island. We included these samples in our analysis. *Anaplasma ovis* was detected in one, *A. bovis* in the second, a co-infection *A. ovis*/*A. bovis* in the third one and no *Anaplasma* in the fourth one.

### 3.6. *Anaplasma* spp. sequence analysis

#### 3.6.1. Anaplasma sp

One 16S rRNA partial sequence from sheep (512 bp) did not align in a BLASTn search with any of the *Anaplasma* species we aimed at with the multiplex *gltA*-based nested PCR. This sequence has 100% identity with *A. platys* detected in *Rhipicephalus* spp. collected on Spanish Ibex (*Capra pyrenaica*) in Spain (PP530072) and an *Anaplasma* sp. from zebra and leopard in South Africa (OQ909488), but is also highly similar to a wide range of different candidatus *Anaplasma* species. Alignement with about 90 *A. platys* 16S rRNA sequences from diverse origins (dogs, ticks, other) highlighted the presence of 7 SNPs specific to both our sequence and the sequence from Ibex in Spain (data not shown), casting doubts on the *A. platys* species identity. We therefore referred to *Anaplasma* sp. for this sequence (accession number LC897980). The phylogenetic analysis indeed placed this isolate in a clade separate from *A. platys* from dogs from diverse countries and among various *A. platys*-like sequences (Fig. 7). This grouping was supported by a 86 % bootstrap value.

**Fig. 7.**
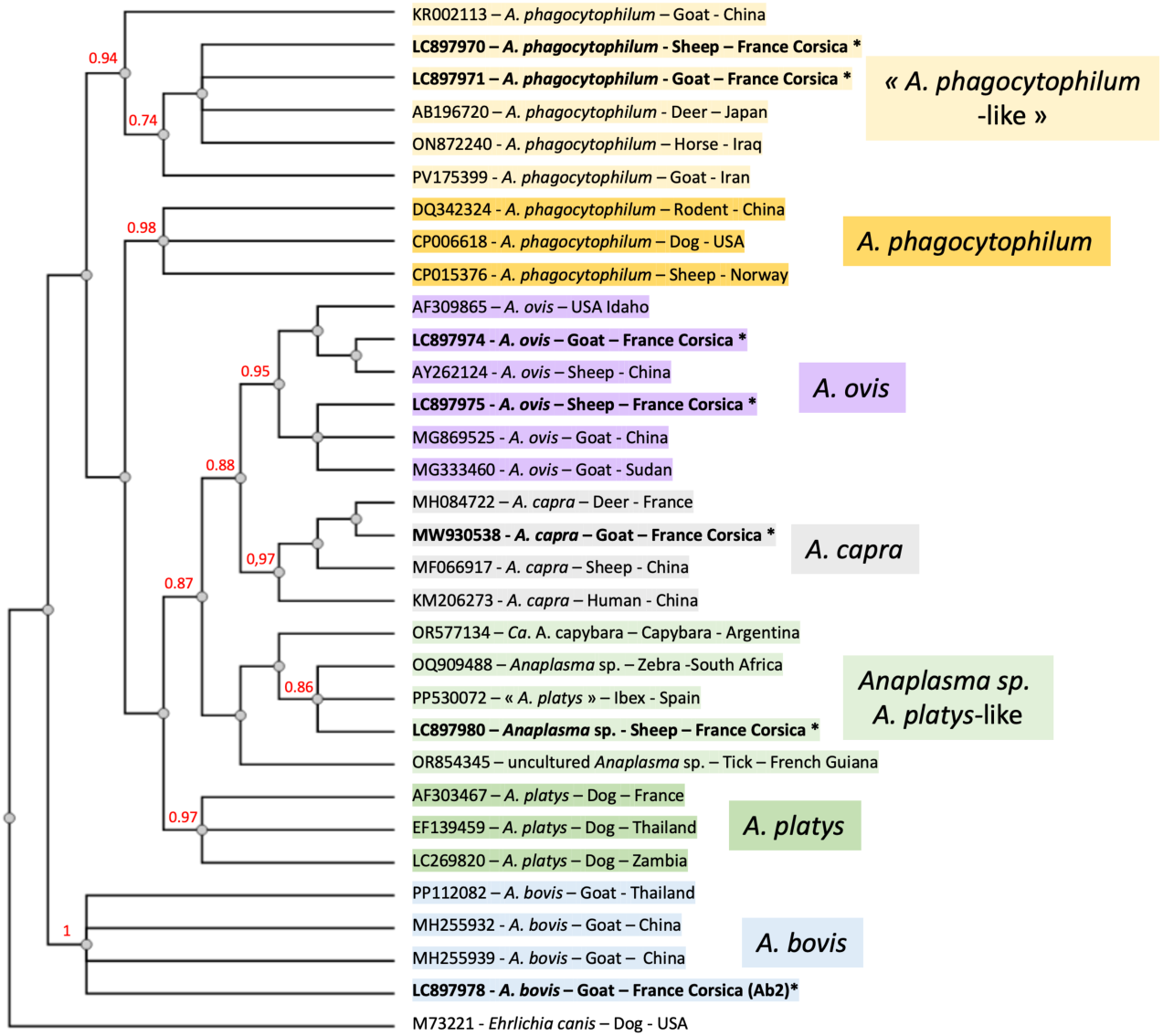
Phylogenetic tree comparing *Anaplasma* spp., based on 501 bp of 16 S rRNA sequences from this study (in **bold with an \***) with isolates representative of the diversity of species, hosts and geographical origins. An alignment with MAFFT followed by BMGE cleaning resulted in all 501 bp being kept. The tree was constructed by the Maximum Likelihood method implemented in the FastTree algorithm with 1000 bootstraps, on the platform http://ngphylogeny.fr. The GTR evolutionary model was chosen, with a gamma distribution of substitution rates. Seven sequences from this study were closely related with five species or species complexes: *Anaplasma phagocytophilum*-like sequences that were distinct from classical *A. phagocytophilum*; *A. ovis*; *A. capra*; *A. platys*-like and *A. bovis*. *Ehrlichia canis* (M73221) was used as an outgroup. Bootstrap values over 70 are shown.

#### 3.6.2. *Anaplasma phagocytophilum* and *A. phagoctophylum-*like

Six partial *A. phagocytophilum* 16S rRNA sequences were obtained, one from goat and 5 from sheep. Five of these sequences were identical, two representative sequences one from sheep (accession number LC897970) and one from goat were deposited (accession number LC897971). One shorter sequence from sheep differed at two positions (accession number LC897972). The longest sequence is also 100% identical to *A. phagocytophilum* from horse in Iraq (ON872235) and from deer in Japan (AB196720), close to *A. phagocytophilum* from goat in Iran (PV175399, 99.81%, one SNP) and China (KR002113, 99.43%, three SNPs) but rather different from the 16S rRNA sequence from sheep in Norway (CP015376, 98.7%, seven SNPs). The phylogenetic analysis indeed placed these isolates in a clade significantly separate from *A. phagocytophilum* from dog, sheep (complete genomes) and rodent from diverse countries in the world and should probably be considered as *A. phagocytophilum*-like (Fig. 7). The separate cluster of *A. phagocytophilum*-like sequences was supported by a boostrap value of 94 %.

One *A. phagocytophilum* partial *gltA* sequence (541 bp) was obtained from a goat (accession number LC902573). This sequence has a high similarity with *A. phagocytophilum gltA* sequences from human (AF304138) and sheep from Norway (CP015376, 99.82%, one SNP), human from Slovenia (CP166491), dog (MK804079) and rodent (*Rattus tanezumi*, MK804080) from South Africa (99.63%, two SNPs).

At least two to three variants of true *A. phagocytophilum* and *A. phagocytophilum*-like isolates are probably circulating in the small ruminants’ population in Corsica, and the most frequent one might be undetected using our designed primers as we failed to amplify *A. phagocytophilum* partial *gltA* gene in the six animals from which 16S rRNA sequences were obtained.

#### 3.6.2. Anaplasma ovis

*Anaplasma ovis* partial 16S rRNA sequences (302 to 525 bp) were obtained from 35 animals: 12 asymptomatic sheep, 22 asymptomatic goats and one symptomatic goat. They differ at 4 positions, three of them being unique and different between sequences, and could be due to amplification or sequencing errors. On the contrary, the fourth SNP split the 29 longest sequences into two groups statistically linked to the host of origin (Fisher exact test, p = 0.0002). One group Ao1goat was more abundant in goat compared to sheep (19 goat and 2 sheep) and the second group Ao2sheep was more abundant in sheep compared to goat (7 sheep and one goat). The two types of sequences were detected in one goat farm (C3). The Ao2sheep sequences were detected in 4 sheep farms (O1, O2, O5 and O6) while the Ao1goat type sequences were detected in two other sheep farms (O7 and O10). Representative sequences of each group from each host were deposited under accession numbers LC897973-76. The similarity searches by blast were not relevant due to the high conservation of the 16S rRNA sequence in the amplified region. Indeed, the Ao2sheep sequences (LC897973-74) were identical to *A. ovis* (KX579068, CP015994). The Ao1goat type sequences (LC897975-76) were identical to more than hundred sequences mainly from uncultured *Anaplasma* sp. (OP020290), diverse *A. marginale* from cattle in Africa (KU686789, PV133719), or *A. ovis* from goat in Sudan (MG333460) and China (MG869525). In the phylogenetic analysis, both groups cluster within *A. ovis* clade (Fig. 7).

The 19 partial *gltA* sequences obtained for *A. ovis* (213 to 338 bp) originated from 10 different farms, mainly from asymptomatic infected sheep (13 sequences) but also from infected goat (6 sequences, 2 from symptomatic and 4 from asymptomatic animals) and were identical, whatever the 16S rRNA sequence group (Ao1goat or Ao2sheep). These sequences are also identical in the amplified region to *gltA* sequences of *A. ovis* from sheep or ticks in China (KX579068, PP117099, OM648114), sheep in Bosnia and Herzegovina (PP354011) and Tunisia (MN238923), but also from goat (MN238922) and dromedary (*Camelus dromedarius*) from Tunisia (MN238937). Representative sequences obtained in this study were deposited (accession number LC902574 for *A. ovis* from sheep and LC902575 for *A. ovis* from goat).

#### 3.6.2. Anaplasma capra

Nine partial *gltA* sequences (465 to 527 bp) of *A. capra* were obtained from eight asymptomatic infected sheep and one goat. They were identical to the previously published sequences amplified from the same farms (MW930533 and MW930535) and belong to *A. capra* clade II and were therefore not deposited. The partial 16SrRNA sequences were also already deposited (MW930536-38). As already published, the 16SrRNA sequences group with other *A. capra* sequences (Fig. 7).

Confirmations of co-infections were obtained from partial *gltA* sequences from singleplex nPCR products of co-infecting *A. capra* and *A. ovis* from five co-infected animals (four sheep and one goat), as well as for co-infecting *A. ovis* and *A. bovis* from one sheep. The identities from the three co-infecting species (*A. capra*, *A. ovis* and *A. bovis*) present in one asymptomatic goat were also confirmed.

#### 3.6.2. Anaplasma bovis

Six amplicons generated using *A. bovis gltA*-specific primers were sequenced. This allowed us to confirm amplification specificity and gather information on the genetic diversity of the circulating isolates. The six sequences, ranging from 501 to 592 bp in length, obtained from *A. bovis*-positive caprine (4 samples) or ovine (2 samples) blood samples, each originating from different geographic locations.

These sequences clustered into two distinct groups, designated Ab1 (*Anaplasma bovis* 1) and Ab2 (*Anaplasma bovis* 2). The three Ab1 sequences were identical and were detected in two asymptomatic sheep (OVI20-07-E1 and OVI20-10-E5) from different flocks, and one asymptomatic goat (CAP20-08-E11). The three Ab2 sequences originated from goats; two were symptomatic (CAP20-02 and CAP20-03) and one was asymptomatic (CAP20-06-E11). The Ab2 *A. bovis* sequences from the two symptomatic goats were identical, while the sequence from the asymptomatic goat differed by two nucleotides over 592 bp, showing 99.7% identity. A sequence identity of 87.4% was observed between the Ab1 and Ab2 types. Notably, Ab1 and Ab2 sequence types could co-circulate within the same goat herd. Sequences of Ab1 from asymptomatic sheep and goat (respectively LC902576-77 and LC902578) and Ab2 from symptomatic and asymptomatic goat (respectively LC902579-80 and LC902581) were deposited.

The nucleotide similarity index was determined by comparing our sequences with representatives of the seven clades of *A. bovis gltA* sequences recently described by Aung et al., 2024 (Table 4). Sequences of Ab1 and Ab2 were different from any of these clades. Ab1 sequences showed higher similarity (89% to 95% identity) to any of the established *A. bovis* clades compared to Ab2 sequences (86% to 89% identity). Furthermore, sequence identities between our Ab1 or Ab2 sequences and the American *A. bovis*-like strains were lower (78% and 75%, respectively) than the identities observed with the various *A. bovis* clades (86% to 95%).

**Table 4.**
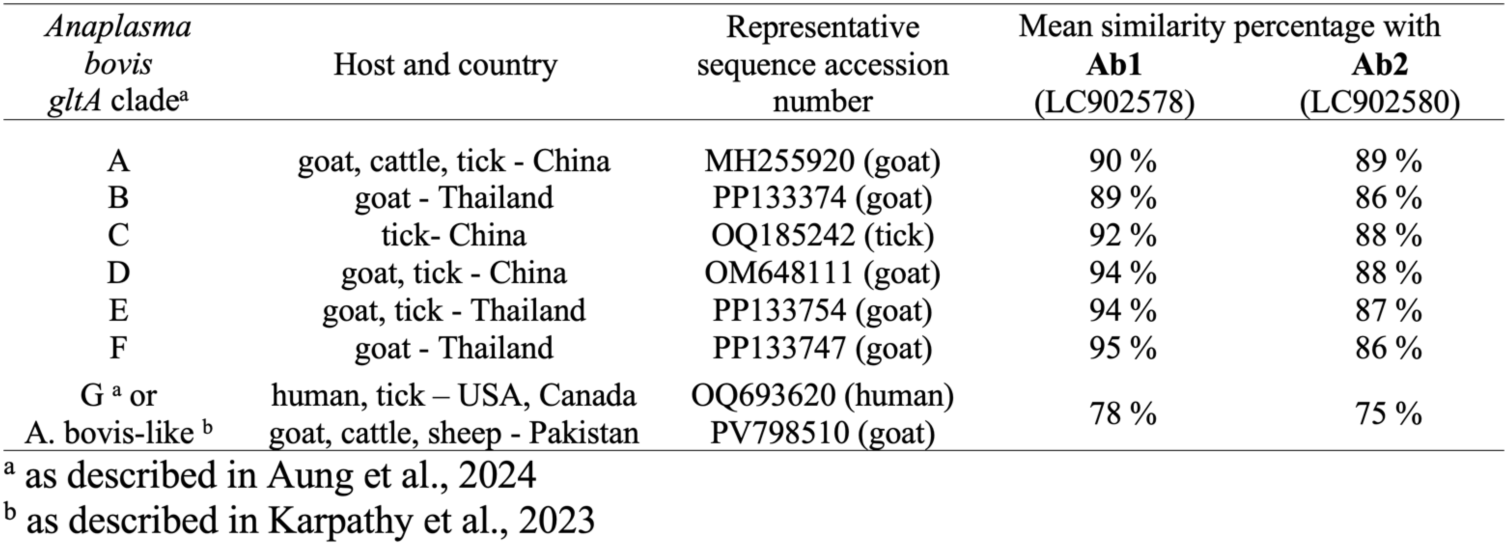
Nucleotide similarity index calculated between a *gltA* representative sequence of each *Anaplasma bovis* (Ab) type from this study and representative *gltA* sequences from the seven described *A. bovis* clades ^a,b^.

The phylogenetic analysis confirmed the position of Ab1 and Ab2 in two separate clades different from all other six already defined *A. bovis gltA* clades (Fig. 8). While the Ab1 clade is closely related to the E and F clades within the well-supported group containing the A to F clades, the Ab2 clade was found to be well-separated from this main group of seven *A. bovis gltA* clades.

**Fig. 8.**
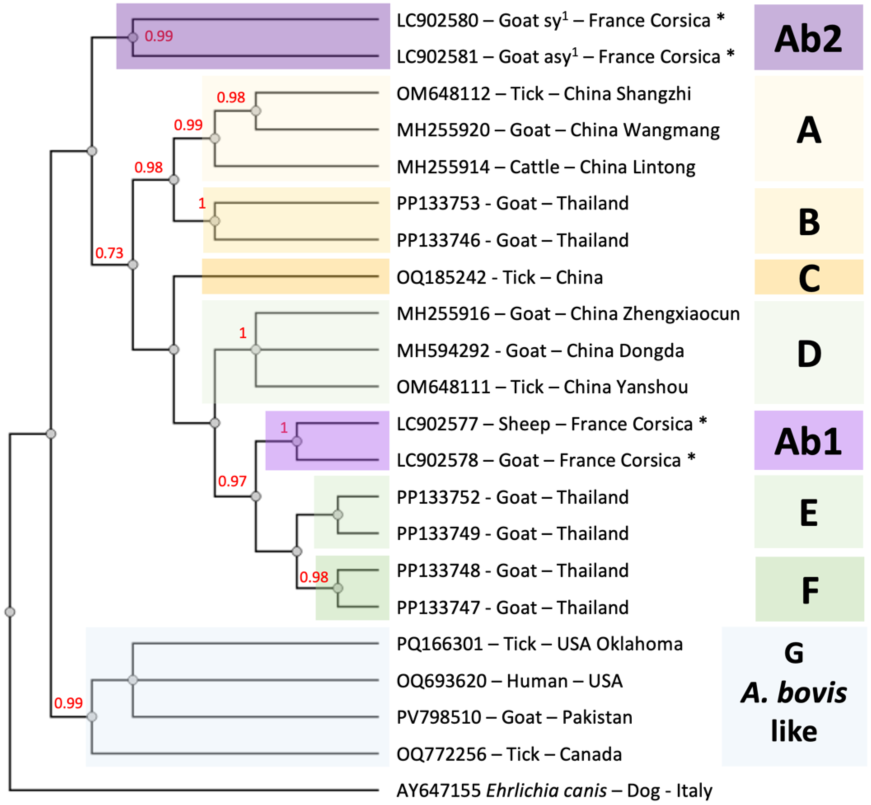
Phylogenetic tree of *Anaplasma bovis* and *A. bovis-*like sequences, based on 416 informative base pairs, after alignment with MAFFT software and curation with the BMGE method. Sequences from this study (with an *, ^1^: sy = symptomatic; asy = asymptomatic) were compared with representatives of the 6 clades A to G described by Aung et al. (2024). The tree was constructed by the Maximum Likelihood method implemented in the FastTree algorithm with 1000 bootstraps, on the platform http://ngphylogeny.fr. The GTR evolutionary model was chosen, with a gamma distribution of substitution rates. Sequences from the Ab1 cluster were closely related to the E and F clades, sequences from the Ab2 cluster stood apart from known clades. Bootstrap values over 70 are shown.

On the 138 comparable amino acids positions 44 positions were found to be variable between the 7 published GltA clades (including the A. bovis-like clade G) and the Ab1 and Ab2 sequences (Fig. 9). The two clades from this study differed at 16 positions (88.4% identity). Half of these modifications (8/16) were unique to the Ab2 sequence when compared to all the other clades, including the Ab1 group. By comparison, only one amino acid modification is unique to either Ab1, A, and F clades, three to either clade B or C and none for clades D and E. The *A. bovis*-like or clade G sequence is the most different, with 24 unique amino acid modifications compared to the eight *A. bovis gltA* clades.

**Fig. 9.**
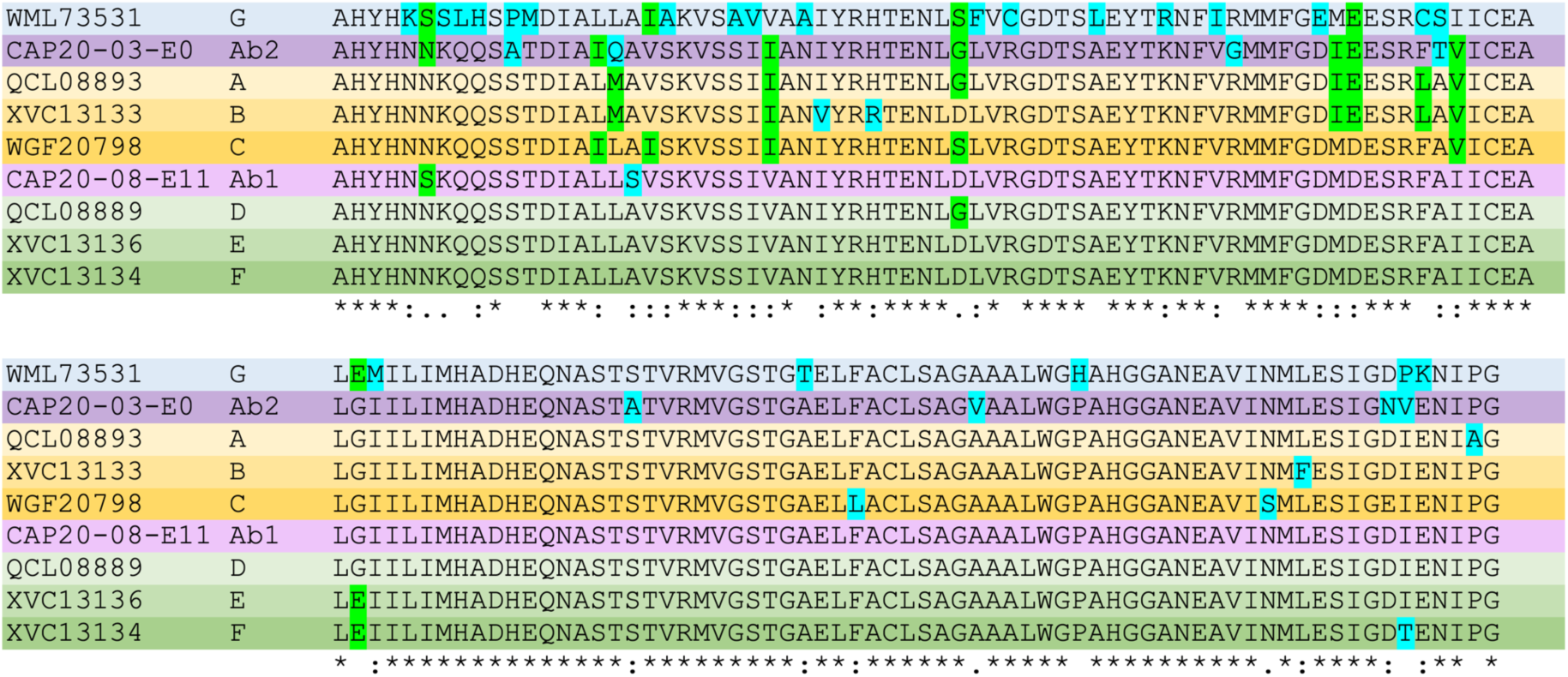
Amino acids alignments of representative GltA partial sequences from all *A. bovis* clades and the Ab1 and Ab2 groups from this study. The clade names as well as the background colors of each clade correspond to those used in Aung et al., 2024 and in Fig. 8. WML73531 (clade G) represents the *A. bovis*-like sequences. AA unique to the clade. AA variable and shared between clades.

To confirm the species identity of the detected isolates, 16S rRNA gene sequencing was performed, yielding six identical sequences, from one sheep (accession number LC897979) and five goats symptomatic (representative sequence accession number LC897977) or asymptomatic (representative sequence accession number LC897978). Three of these sequences originated from the same animals that provided the Ab2 *gltA* sequences previously described (accession numbers LC902580 and LC902581). For the remaining three, corresponding *gltA* sequences were not obtained, making it impossible to determine if they belonged to the *gltA* Ab1 or Ab2 *A. bovis* types. In the phylogenetic analysis, the 16S rRNA sequence clustered with other *A. bovis* sequences in a well-supported clade (Fig. 7).

We then compared these six 16S rRNA gene sequences with available 16S rRNA sequences from isolates whose *gltA* sequences were classified by Aung et al. (2024). No differences in the 16S rRNA sequences between the isolates from the current study and the 16S rRNA sequences of *gltA* clade A (OM569659, OM569660, MH255935, MH255941) or *gltA* clade D (MH255926, MH255932, OM569658, OQ1325531, OQ132533) were observed. In the same 16S rRNA region, two nucleotide differences were detected when compared to the American *A. bovis*-like sequences (data not shown).

## 4. Discussion

In this study, we developed a *gltA*-based multiplex nested PCR (nPCR) assay to detect the five most common *Anaplasma* species in small ruminants: *A. ovis*, *A. capra*, *A. phagocytophilum*, *A. bovis*, and *A. marginale*. After confirming the primers’ specificity, we applied the assay to a cohort of sheep and goats from Corsica, France, which had been previously tested for *A. capra*. Our results confirmed the presence of *A. ovis* and *A. capra* and, for the first time, provided evidence of *A. phagocytophilum* and *A. bovis* circulating in small ruminants on the island. Additionally, a control 16S rRNA nPCR assay enabled us to report the first detection of a probably new species of *Anaplasma* in Corsica.

The singleplex, *gltA*-based PCRs were generally more efficient at detecting infections compared to the duplex or multiplex nPCR protocols. For *A. ovis*, *A. capra*, and *A. bovis*, the efficiency of the singleplex PCRs was comparable to or higher than our standard 16S rRNA nested PCR (nPCR) protocol. This demonstrates that using three separate singleplex PCRs, these three species can be reliably detected and identified, even in a co-infected blood sample. This approach achieves an efficiency similar to that of a multi-copy gene target nPCR protocol, but without the need for sequencing.

For these species, the duplex PCR and multiplex nPCR showed a comparable (for *A. capra*) or lower (for *A. ovis* and *A. bovis*) efficiency than their respective singleplex protocols. This reduced performance is likely due to competition between primers during amplification and, in the case of the nested protocol, the preferential amplification of one target over others during the initial PCR.

The singleplex PCR designed to detect *A. phagocytophilum* proved to be inefficient to detect the circulating genetic variants. We hypothesize this could be linked to the well-documented genetic diversity of *A. phagocytophilum* (Scharf et al., 2011), even at a local scale (Jouglin et al., 2017). This diversity might have resulted in sequence heterogeneity within the *gltA* gene, which would prevent our primers from binding correctly. In Corsica, at least two genetic variants seem to circulate, and we hypothesize that the main variant detected by 16S rRNA amplification might have a different *gltA* sequence not amplified with the designed primers.

An *a posteriori* analysis of *gltA* sequences deposited since our primer design (OP585591-602 and OM6481119) supports this possibility. This analysis revealed significant polymorphism in both the PCR1 forward primer sequence (5 nucleotides) and the PCR2 *A. phagocytophilum*-specific forward primer sequence (11 nucleotides). While these sequences originate from distant locations (China and Chile), they demonstrate that the regions initially selected for primer design—which appeared conserved at the time—can in fact vary. Therefore, we cannot exclude the possibility that undetected variants were circulating in our cohort, which would only have been captured by the more universal *Anaplasma* spp. 16S rRNA nPCR.

Combining the different PCR protocols, we found a high infection rate of *A. ovis* in both goats (89.5%) and sheep (64.3%). Our findings are consistent with previous studies in Corsica that also demonstrated high prevalence rates, such as 42.3% in sheep (Dahmani et al., 2017) and 52% in goats (Cabezas-Cruz et al., 2019). In contrast, one study reported a low infection rate in sheep (2.7%, 2/74 sheep), limited to only one of the three herds sampled (Defaye et al., 2022). This discrepancy could be linked to the detection method used (PCR, qPCR, nPCR, or microfluidic real-time PCR), as all these studies were conducted in the same general areas of Corsica within a limited timeframe (2014-2020).

The high prevalence of *A. ovis* in Corsica reflects its status as a highly frequent and endemic pathogen in small ruminants across the Mediterranean basin. For instance, prevalence rates have been reported at 82.5% in sheep in Portugal (Renneker et al., 2013), 77.2% in sheep on the island of Lesvos in Greece (Saratsis et al., 2022), and ranging from 7-87% in sheep and 19% in goats in Italy (De la Fuente et al., 2005; Torina et al., 2008). Other examples include Türkiye (69.4% in sheep and 44.7-46.1% in goats; Ulucesme et al., 2023; Karatepe et al., 2025), Tunisia (35.6-93.8% in sheep and 46-70.3% in goats; Ben Said et al., 2015; Belkahia et al., 2017; M’ghirbi et al., 2022), and Algeria (61.7% in sheep and 54.2% in goats; Aouadi et al., 2017; Chadi et al., 2024). In Spain, 69 out of 70 sheep herds were found to be infected (Ruiz et al., 2025). The impact of *A. ovis* on animal health is difficult to quantify, as most studies are conducted on asymptomatic animals. Furthermore, comparative studies on both poor health and asymptomatic animals have revealed similar prevalence rates (37% and 47.3%, respectively; Torina et al., 2010; Torina and Carapacca, 2012), suggesting that the pathogen’s clinical effect may not be straightforward.

We detected a 16S rRNA polymorphism in the *A. ovis* isolates that was statistically linked to the host of origin (sheep or goat). While this host-related divergence was not observed in the *gltA* partial sequences, it is important to note that only short fragments of *gltA* were obtained. A previous study examining *msp4* polymorphism in the Corsican goat population found one dominant variant, COR1, in 94.6% of infected goats, with the remaining five variants detected only once or twice (Cabezas-Cruz et al., 2019). Since the Ao1goat 16S rRNA variant similarly accounts for 95% of the *A. ovis* goat isolates in our study, it would be highly valuable to determine if a correspondence exists between this Ao1goat variant and the *msp4* COR1 variant.

*Anaplasma capra* has been previously detected in the same small ruminant population in Corsica (Jouglin et al., 2022). Our findings are consistent with this earlier work, though with slight differences: the present study detected *A. capra* in one additional sheep flock (O6, with one infected sheep) and one more animal in flock O5, compared to the previous study which used different primers. Our results show *A. capra* infection rates of 34.3% in sheep and 13.2% in goats, with a higher number of infected sheep farms (5/7) compared to goat farms (1/4). In contrast, another study in Corsica (Defaye et al., 2022) reported an extremely low infection rate of 1.4% in sheep (1/74). First discovered in goats in China approximately 13 years ago (Li et al., 2015), *A. capra* is now widespread in small ruminants across Asia (Lin et al., 2023; Altay et al., 2024). In Europe, it was first identified in deer in mainland France (Jouglin et al., 2019) and has since been found in small ruminants in other Mediterranean countries (Greece, Türkiye), in cattle (Morocco, Türkiye), and in deer (Spain). However, reported infection rates in these regions are generally lower than those we found in Corsica (Elhachimi et al., 2021; Remesar et al., 2022; Altay et al., 2022; Saratsis et al., 2022; Oguz et al., 2024).

In Mediterranean countries, *A. phagocytophilum* has been detected with low infection rates in sheep in Italy (3%; Torina et al., 2008) and in both sheep (0.7%) and goats (1.7%) in Türkiye (Aktas et al., 2021). It has also been reported in goats in Cyprus (Chochlakis et al., 2009) but not in northern Tunisia (Belkahia et al., 2017). Our results are consistent with these data, with low infections rates of 7.1 % in sheep and 5.3 % in goats. *Ixodes ricinus*, the main vector of *A. phagocytophilum* in Europe, is not frequent in Mediterranean regions (Gray et al., 2024). In Corsica, *I. ricinus* represents only 5.7% of the ticks collected from cattle and has never been collected from sheep or goats on this island (Grech-Angelini et al., 2016; Cabezas-Cruz et al., 2019), indicating a low abundance on these hosts, and subsequently a low transmission.

We did not detect *A. marginale* in our study, despite its confirmed presence in Corsica in cattle and in *Rhipicephalus bursa* ticks collected from sheep and goats (Cicculi et al., 2019; Grech-Angelini et al., 2020). The presence of *A. marginale* in small ruminants has been documented in other regions, including in sheep (3.3%) and goats (18.8%) in Italy (Torina et al., 2008), in sheep in Algeria (Chadi et al., 2024), and in *R. bursa* collected from sheep or goats in Portugal (Ferrolho et al., 2016). Based on these findings, we hypothesize that the prevalence of *A. marginale* in small ruminants in Corsica may be too low to have been detected within our study population.

We report the presence of a probably new *Anaplasma* species for the first time in Corsica, in only one sheep (0.9%). This 16S rRNA sequence has 100% identity with an *Anaplasma* named *A. platys* detected in *Rhipicephalus* spp. collected on Spanish Ibex (*Capra pyrenaica*) in Spain (Masià-Castillo et al, 2025) and an *Anaplasma* sp. (ST_KNP_5) from zebra and leopard in South Africa (Makgabo et al., 2023). The sequence groups with several *Anaplasma* species (*A. platys*) or candidatus species (candidatus *A. camelii, candidatus A. capybara*) with high sequence identities, and further investigations are required, including the analysis of other genes for the discrimination of species within this genus. We detected this probably new species only with the 16S rRNA amplification, due to the species specificity of the *gltA* primers used.

We characterized *Anaplasma bovis* infection for the first time in Corsica, with a 4.3% infection rate in sheep and a significantly higher rate of 18.4% in goats. In other Mediterranean countries, this species has been found in Tunisia with varying infection rates (0-42.7% in sheep and 0-23.8% in goats, depending on the study; Belkahia et al., 2017; Ben Said et al., 2015; M’ghirbi et al., 2022), in sheep in Greece (1.6%; Saratsis et al., 2022), and in sheep from Sardinia, Italy (Zobba et al., 2014). Interestingly, it has not been detected in small ruminants in Algeria (Chadi et al., 2024) or Türkiye (Karatepe et al., 2025).

Our study also identified two new *A. bovis gltA* clades, which are distinct from the six clades previously described in isolates from China and Thailand (Lu et al., 2023; Aung et al., 2024; Tang et al., 2024). These are the first *A. bovis* sequences from European isolates to be described. While variants from three different clades circulate in China and Thailand, we found two other clades circulating in Corsica, suggesting a pattern of geographical clustering. Further research on European and other continental isolates may help us better understand the spatial structure of *A. bovis* and its potential link to vector species distribution (Huang et al., 2025).

The *gltA* Ab2 variants we found differ greatly from the variants of the seven other clades, especially at the amino acid level, even though their 16S rRNA sequences are 100% identical to other *A. bovis* sequences. Interestingly, we observed that symptomatic goats were infected with this new variant, which may suggest that it has a different virulence profile than other variants.

In our study, *A. bovis* was never detected as a mono-infection in asymptomatic animals; it was always found as a co-infection with *A. ovis,* except for one symptomatic goat. This trend has also been observed in Greece (Saratsis et al., 2022) and, more significantly, in Tunisia (Belkahia et al., 2017). Co-infections with two *Anaplasma* species were frequent in both sheep (35.1%) and goats (32.4%). As expected, most of these co-infections involved the two most common species, *A. ovis* and *A. capra* (54.8%). Co-infections with three species - *A. ovis*, *A. bovis*, and *A. phagocytophilum* - have also been documented in sheep and goats in China (Zhang et al., 2016). The ability of *A. ovis* to establish long-lasting infections (4–6 years) in sheep (Ruiz et al., 2024) may increase the likelihood of a secondary infection with less common species over time.

In Corsica, *R. bursa* is the dominant tick species on small ruminants, accounting for over 90% of all collected ticks (Grech-Angelini et al., 2016, 2020; Cabezas-Cruz et al., 2019; Cicculi et al., 2019). While *R. bursa* is considered the primary vector of *A. ovis*, its role in transmitting the other four *Anaplasma* species remains unclear. Other potential vectors are not frequently found on small ruminants in Corsica. For example, *Ixodes ricinus*, the main vector for *A. phagocytophilum* in Europe (Gray et al., 2024), is not common in our study area. Similarly, while *Haemaphysalis* species (*H. longicornis* in Asia and *H. concinna* in France) are suspected vectors of *A. capra* (Altay et al., 2022; Jouglin et al., 2025), they are not abundant on the sampled hosts in Corsica. The vectorial competence of several suspected vectors of *A. bovis* also remains to be determined (Atif, 2016; Rar et al., 2021).

## 5. Conclusion

Ovine and caprine populations in Corsica show high rates of infection and co-infection with various *Anaplasma* species, primarily *A. ovis* and *A. capra*. The discovery of *A. bovis* in Corsica, exclusively as a co-infecting species and featuring novel genetic variants, warrants close attention. Notably, one of these new variants was detected in two out of four symptomatic goats, in samples submitted to our laboratory for confirmation of the presence of *Anaplasma*, justifying further clinical and epidemiological investigation. While a previous study found no link between *A. ovis* infection and health indicators in apparently healthy goat flocks (Cabezas-Cruz et al., 2019), co-infections were not assessed, highlighting a critical gap. Finally, given that *Rhipicephalus bursa* is the dominant tick species found on small ruminants in Corsica, future studies must clarify its central role in the vectorial competence and transmission routes of these bacterial species.

## CRediT Taxonomy

**MJ**: Investigation - Methodology - Project Administration - Supervision - Validation - Visualization - Writing - Original Draft - Writing - Review & Editing

**LR:** Investigation - Writing - Review & Editing

**CB:** Supervision- Writing - Review & Editing

**SB**: Data Curation - Formal Analysis - Visualization - Writing - Review & Editing

**LM**: Conceptualization - Formal Analysis - Funding Acquisition - Methodology - Project Administration - Supervision - Validation - Visualization - Writing - Original Draft - Writing - Review & Editing

## Acknowledgements

We would like to thank Professor Furhan Iqbal (Bahauddin Zakariya University, Pakistan) for donating DNA from *A. marginale* infected sheep.

## Declaration of generative AI and AI-assisted technologies in the writing process

During the preparation of this work, LM used Gemini in order to improve the English language. After using this tool, the authors reviewed and edited the content as needed and take full responsibility for the content of the publication.

## Data availability

The datasets used and/or analyzed during the current study are available from L.M. on reasonable request.

